# Soil bacteria may be the natural reservoirs of drug resistance genes

**DOI:** 10.1101/2024.09.06.611735

**Authors:** Salman Ahmad, Asad Bashir Awan, Sofia Irfan, Abdul Haque

## Abstract

Soil bacteria are the main source of antibiotics because they produce them naturally to get territorial advantage. We collected deep soil samples and characterized cultivable microbes. We used morphological and cultural characters, biochemical reactions, 16s RNA PCR and DNA sequencing to identify the isolates. The isolates included *Pseudomonas* spp (12), *Shigella* spp (2), *E. coli* (1), *Klebsiella/Citrobacter* spp (1), *Micrococcus spp* (2), unidentified *Bacillus* spp.(18), *Bacillus paramycoides* (3), *Paenibacillus lautus* (2), *Bacillus pacificus* (2), and *Lysinibacillus pakistanensis* (1). Out of these 44 isolates, 33 (75%) were multi-drug resistant. Clinically relevant and clinically irrelevant bacteria had similar drug resistance patterns (88.9% and 88.0%, 88.9% and 88.0%, 100 and 96.0%, 61.1% and 92%, 83.3% and 96%, 33.3% and 36%, 55.6% and 68.0%, and 83.3% and 60%) towards Ampicillin, Amoxicillin, Oxacillin, Azithromycin, Streptomycin, Gentamicin, Ceftriaxone and Sulfamethoxazole respectively. The observation that bacteria which cannot colonize humans/animals and therefore cannot enter the horizontal drug resistance gene transfer cycle in clinical settings also have a large and similar arsenal of drug resistance genes, may indicate that they are actually the natural reservoirs. Because they are more dynamic in their ability to survive in different conditions, they provide better fitness for maintenance of these reservoirs.

**Impact Statement:** We think this study has raised interesting questions and further probing at a larger scale and a greater depth will provide new insight that may be very helpful for understanding the phenomenon of spread of drug resistance among bacteria which is one of the paramount challenges for humanity.

## Introduction

It has been noticed that resistance against antibiotics is common among soil bacteria.^1^ Soil bacteria produce antibiotics as a result of a competitive mechanism for survival. Genes are transferred from one bacterium to other by horizontal gene transfer.^2^

In 21^st^ century, the major global challenge is drug-resistant pathogenic bacteria that pose risks to public health; and most of the resistance genes found in these bacteria may be of soil origin. ^3^ Resistance carrying genes have been found in the cultivable bacteria isolated from a cave by the anthropogenic influence for 4 million years or in a DNA isolated from 30,000 years old permafrost sediment.^4^

Soil contamination is also a factor playing an important role in emerging bacterial drug resistance. Xenobiotics which are synthesized chemically, are recalcitrant to biodegradation. These compounds released by pharmaceutical and metal industries, and fertilizer manufacturing units challenge the bacteria to adopt to severe conditions for growth, owing to their ill effects on soil texture and nutritional values.^5^

The purpose of this study was to characterize (including drug resistance patterns) cultivatable bacteria from soil samples with the purpose to compare the data of clinically relevant and clinically irrelevant isolates to get an insight into the mechanisms of horizontal transfer of drug resistance genes among these bacteria.

## MATERIALS & METHODS

### Collection of soil samples

Eleven soil samples were collected from different regions of Pakistan; 4 from Punjab, 3 from Khyber Pakhtunkhawa, 2 from Baluchistan and 2 from Gilgit Baltistan. Samples were collected from locations where human and animal movement was infrequent. Soil samples were taken at a depth of six inches to avoid surface contamination. Precautionary measures were observed during sample collection such wearing gloves to avoid contamination by commensal bacteria on human body. Sample size was roughly 200 grams. Samples were tightly closed and stored in plastic zipper bags at 4^°^C.

### Isolation & selection of drug-resistant bacteria

The collected soil samples were centrifuged after addition of equal volume of dH_2_O (20g in 20ml). After centrifugation at 10,000 rpm for 15 minutes, 1 ml supernatant was taken and spread onto a Mueller Hinton Agar plate and paper sensitivity discs (Oxoid, UK) of 8 commonly used antimicrobial drugs representing 5 different antimicrobial groups were applied. These antimicrobials included Ampicillin (10µg), Amoxycillin(10µg), Oxacillin(5µg), Azithromycin(15µg), Streptomycin(10µg), Gentamicin(10µg), Sulphamethoxazole (25µg) and Ceftriaxone(30µg). Observations were made after overnight incubation at 37°C. Results were interpreted according to guidelines of Clinical and Laboratory Standards Institute.^6^

Samples were taken from the vicinity of sensitivity discs which had no inhibition effect and sub cultured on Nutrient Agar plates to get pure cultures. Provisional identification was made by Gram staining and biochemical testing by conventional methods.

### PCR-based confirmation of isolates

After the morphological and biochemical identification, the isolates showing antimicrobial resistance were confirmed by PCR. DNA extraction was done by conventional chloroform-isoamyl-alcohol method. The DNA integrity was checked by electrophoresis on 1% agarose gel before storage at 4ºC.

Universal primers 27 F (5’-AGA GTT TGA TYM TGG CTC AG-3’) and 1492R (5’-CGG TTA CCT TGT TAC GAC TT-3’) were used.^7^ Each 100 µL of the reaction mixture contained 10x PCR buffer 10 µL, 25 mM MgCl_2_, 0.7 nM each dNTP, 25 pM each primer, 5 U of Taq DNA polymerase (Fermantas, Maryland, USA), 10 ng of template DNA, and sterile water to make the volume. Thermal cycler conditions were: initial denaturation at 95°C for 5 minutes followed by 35 cycles of: denaturing at 94°C for 1 minute, annealing at 45°C for 1 minute and extension at 72°C for 2 minutes, and a final extension at 72°C for 7 minutes. The amplicons were observed by using 1.5 % agarose gel electrophoresis.

### DNA sequencing

Amplicons which could not be identified were sequenced by 1^st^ Base, Singapore.

## RESULTS

A total of 44 isolates showed drug resistance by disc diffusion method. Gram positive isolates were identified by their morphological characters such as size, shape, spore formation etc. in addition to their cultural characteristics.

Gram negative isolates were identified by inoculation on TSI medium. Nine, three, one and one isolates showed results consistent with *Pseudomonas/Acinetobacter* species, *Proteus/Provedencia* species, E. coli and *Klebsiella* species respectively.

### DNA sequencing

Nineteen isolates could not be identified by morphology, cultivation and biochemical testing so 16S rRNA genes were amplified to get internal sequences for these isolates. DNA from all the isolates produced the desired amplification product of 1492 bps. Sanger sequencing was performed for the identification of these bacterial isolates. After sequencing, sequences were purified by Bio-edit software for blast on NCBI-BLAST to get the best homology sequences. The sequences were confirmed by BLAST.

Unfortunately, BLAST results could identify only 9 isolates to species level. The results showed that 3 were *Bacillus paramycoides*, 2 were *Paenibacillus lautus*, and there was one isolate each representing *Pseudomonas stutzeri, Bacillus pacificus, Shigella flexneri* and *Lysinibacillus pakistanensis*. Ten isolates which were provisionally identified as *Bacillus* by conventional methods could not be further segregated.

In total, there were 18 clinically relevant isolates and 26 clinically irrelevant isolates. The clinically relevant isolates were *Pseudomonas* spp (12), *Shigella* spp (2), *E*.*coli* (1), *Klebsiella/Citrobacter* spp (1), and *Micrococcus spp* (2).

The number of clinically irrelevant isolates was 26. These included 23 *Bacillus* spp.(including 3 *Bacillus paramycoides* and 2 *Bacillus pacificus*), 2 *Paenibacillus lautus*, and 1 *Lysinibacillus pakistanensis*.

### Antibiotic Susceptibility Test by disc diffusion method

The drug resistant soil isolates (n=44) were tested for their susceptibility towards 8 antimicrobial drugs belonging to 5 different groups. Results were interpreted according to CLSI guidelines ^6^.

Thirty-three (75%) isolates showed resistance to at least 3 antimicrobial drug groups and thus were regarded as MDR (multiple drug resistant). Only 3 (6.8%) of 44 isolates showed resistance towards all the 5 antimicrobial drug groups; 14 (31.81%) were resistant to 4, 16 (36.36%) to 3, 6 (13.6%) to 2 and 5 (11.36%) to only 1 antimicrobial drug group. Results are shown in a Table 1.

**Table 1:** Resistance pattern of isolates towards antimicrobial drug groups.

| **Resistant to Antimicrobial groups** | **No of isolates** | **%** |
| --- | --- | --- |
| Resistant to 5 antimicrobial groups | 3 | 6.8% |
| Resistant to 4 antimicrobial groups | 14 | 31.81% |
| Resistant to 3 antimicrobial groups | 16 | 36.36% |
| Resistant to 2 antimicrobial groups | 6 | 13.6% |
| Resistant to 1 antimicrobial group | 5 | 11.36% |

Out of 44 isolates, 42 (95.45%) were resistant to the penicillins, 14 (31.81%) to cephalosporins, 27 (61.36%) to macrolides, 23 (27%) to aminoglycosides and 30 (68.18%) to sulfonamides. The pattern of resistance was: 86.36%, 84.09%, 93.18%, 61.36%, 52.27%, 18.18%, 31.81%, 68.18% towards ampicillin, amoxicillin, oxacillin, azithromycin, streptomycin, gentamicin, ceftriaxone and sulphamethoxazole respectively (Figure 1).

**Figure 1:**
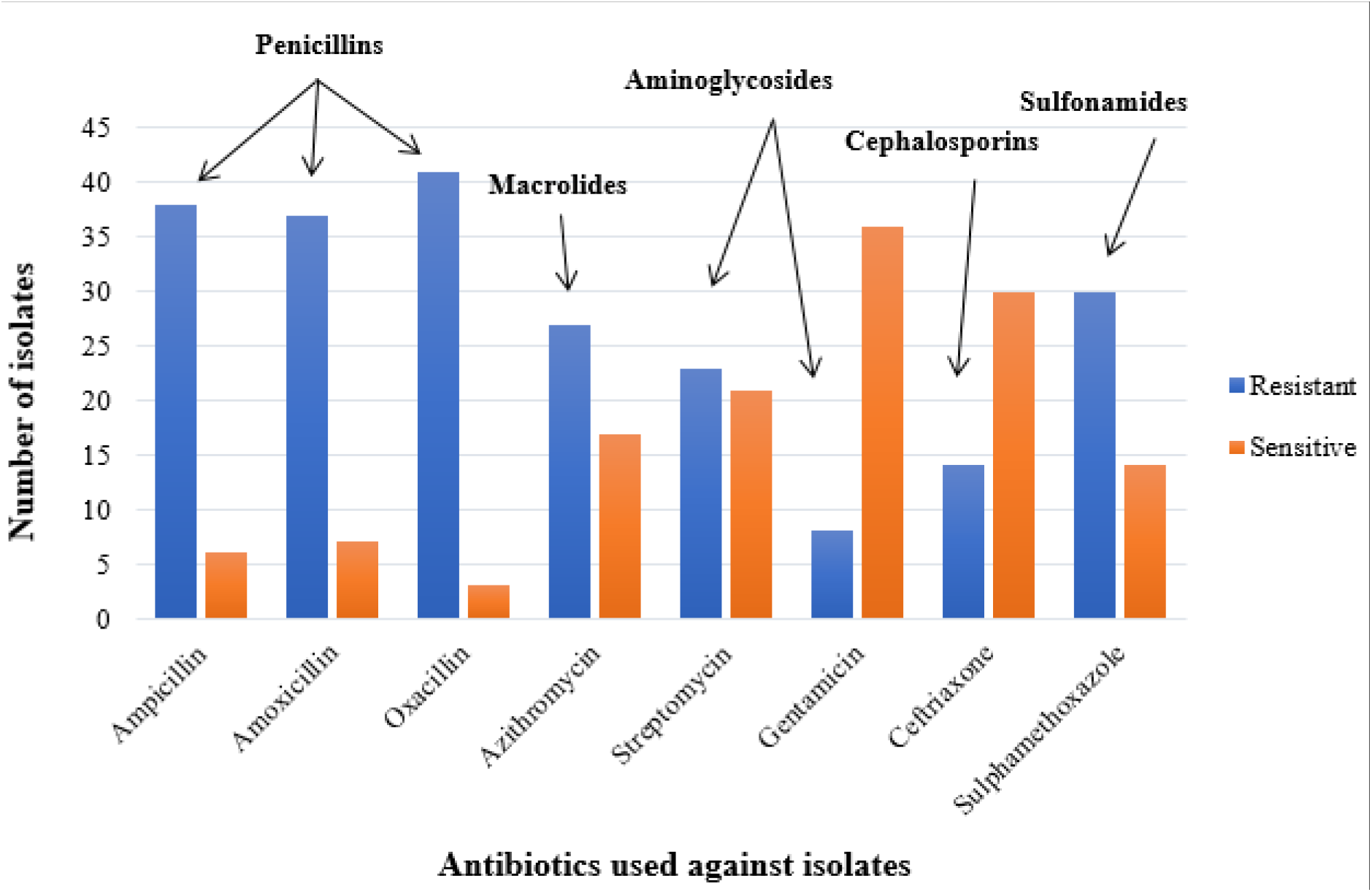
The graphical representation of resistance pattern of the studied isolates against the individual antimicrobial drugs. A total of 42 isolates were tested against the eight antimicrobial drugs belonging to five antimicrobial groups. Among the 42 isolates, 86.36% were resistant to ampicillin, 84.09% to amoxicillin, 93.18% to oxacillin, 61.36% azithromycin, 52.27% to streptomycin, 18.18% to gentamicin, 31.81% to ceftriaxone and 68.18% to sulphamethoxazole.

### Drug resistance pattern of clinically relevant and irrelevant microbes

The results are shown in Table 2. The *Pseudomonas* spp (12), *Shigella* spp (2), *E*.*coli* (1), *Klebsiella/Citrobacter* spp (1), and *Micrococcus spp* (2) were considered as clinically relevant.

**Table 2:** Comparison of drug resistance pattern of clinically relevant and irrelevant microbes.

| Sr # | Antimicrobial | Clinically relevant microbes* | Clinically irrelevant microbes** |
| --- | --- | --- | --- |
| 1 | Ampicillin | 88.9 | 88.0 |
| 2 | Amoxicillin | 88.9 | 88.0 |
| 3 | Oxacillin | 100 | 96.0 |
| 4 | Azithromycin | 61.1 | 92.0 |
| 5 | Streptomycin | 83.3 | 96.0 |
| 6 | Gentamicin | 33.3 | 36.0 |
| 7 | Ceftriaxone | 55.6 | 68.0 |
| 8 | Sulfamethoxazole | 83.3 | 60.0 |

- 245 - 246● Clinically important bacteria: *E.coli*, *Klebsiella/Citrobacter* spp, *Shigella* spp, *Micrococci* 247 and *Pseudomonas* spp - 248● Clinically unimportant bacteria: *Bacillus* sp., *Paenibacillus lautus*, and *Lysinibacillus* 249 *pakistanensis*

Those included in clinically irrelevant group were 23 *Bacillus* spp.(including 3 *Bacillus paramycoides* and 2 *Bacillus pacificus*), 2 *Paenibacillus lautus*, and 1 *Lysinibacillus pakistanensis*.

## DISCUSSION

One of the major global challenges for humanity in 21^st^ century is pathogenic drug-resistant bacteria and many of these have soil as reservoir.^3^ However, more than 99% of soil bacteria observed under microscope cannot be cultured in laboratory.^8^

The objective of this study was to collect soil bacteria from different locations in Pakistan, identify them and to determine the drug resistance patterns of clinically relevant and non-relevant bacteria against commonly used antimicrobial drugs to get an insight to their interactions involving exchange of drug resistance genes.

There were 44 isolates from 11 soil samples. The isolates were identified by conventional and molecular methods as detailed above. We sent 16sRNA gene amplification products of 19 unidentified isolates for sequencing, but only 9 were confirmed by BLAST; 3 were *Bacillus paramycoides*, 2 were *Paenibacillus lautus*, 1 was *Lysinibacillus pakistanensis*, 1 was *Shigella flexneri*, 1 was *Pseudomonas stutzeri* and 1 was *Bacillus pacificus*.

High drug resistance was seen in general as out of 44 isolates, 75% were multi-drug resistant; 6.8%, 31.81%, and 36.36% were resistant to 5, 4 and 3 drugs respectively. Among the remaining 25%, 13.6% and 11.36% showed resistance towards 2 and one drug respectively as shown in Table 1. These findings are in line with existing reports.^9^

Very high resistance was seen against penicillins as 86.36%, 84.09% and 93.18% isolates were resistant to Ampicillin, Amoxicillin and Oxacillin. Similar results have been reported previously.^10^ Azithromycin, Streptomycin and Sulfamethoxazole were moderately effective with 61.36%, 68.18% and 52.27% resistant isolates respectively. Most effective drugs were gentamicin and ceftriaxone with only 18.18% and 31.81% resistant isolates respectively.

In total, there were 18 clinically relevant isolates and 26 clinically irrelevant isolates as described in detail in results section.

The drug resistance patterns were similar among clinically relevant and clinically irrelevant bacteria as detailed in Table 2. It shows that the clinically irrelevant bacteria which have not gone through the cycle of human infections still possessed these drug resistance genes in their natural habitat and to the same extent and on similar pattern as clinically relevant bacteria. Does it mean that all these bacteria have natural reservoirs of these genes -- but we focus only on those which are capable of causing disease?

It is common knowledge that the pathogenic bacteria exchange drug resistance genes by horizontal transfer in a conducive atmosphere (e.g., hospitals) or due to close proximity of healthy persons with patients and carriers. But our study, in which we took samples from a depth of 6 inches in uninhabited areas, shows that they are intrinsically equipped with this arsenal which they share with non-pathogenic bacteria present in soil even without exposure in clinical settings.

This raises the possibility that all bacteria have these reservoirs naturally. Even if we argue that the clinically relevant bacteria present in deep soil somehow passed through horizontal exchange cycle of genes, what about other bacteria which do not have the capacity to colonize humans or animals and these included rare species such as *Bacillus paramycoides, Bacillus pacificus, Paenibacillus lautus* and *Lysinibacillus pakistanensis*.

What is their source of drug resistance genes? Are they themselves the source of these genes and clinically relevant bacteria have benefitted from them? This is plausible as these saprophytes have better fitness to survive and flourish in natural environments such as soil whereas clinically relevant bacteria have specialized to survive and grow in human and animal bodies, i.e., their versatility is not comparable with clinically irrelevant bacteria.

## Acknowledgments

The work was fully funded by AKHUWAT Foundation as part of the AKHUWAT-FIRST Under-Graduate Research Program under AKHUWAT Educational Services.

